# Pathway Coessentiality Mapping Reveals Complex II is Required for *de novo* Purine Biosynthesis in Acute Myeloid Leukemia

**DOI:** 10.1101/2025.02.26.640463

**Authors:** Amy E. Stewart, Derek K. Zachman, Pol Castellano-Escuder, Lois M. Kelly, Ben Zolyomi, Michael D.I. Aiduk, Christopher D. Delaney, Ian C. Lock, Claudie Bosc, John Bradley, Shane T. Killarney, Olga R. Ilkayeva, Christopher B. Newgard, Navdeep S. Chandel, Alexandre Puissant, Kris C. Wood, Matthew D. Hirschey

## Abstract

Understanding how cellular pathways interact is crucial for treating complex diseases like cancer, yet our ability to map these connections systematically remains limited. Individual gene-gene interaction studies have provided insights^1,2^, but they miss the emergent properties of pathways working together. To address this challenge, we developed a multi-gene approach to pathway mapping and applied it to CRISPR data from the Cancer Dependency Map^3^. Our analysis of the electron transport chain revealed certain blood cancers, including acute myeloid leukemia (AML), depend on an unexpected link between Complex II and purine metabolism. Through stable isotope metabolomic tracing, we found that Complex II directly supports *de novo* purine biosynthesis and exogenous purines rescue AML from Complex II inhibition. The mechanism involves a metabolic circuit where glutamine provides nitrogen to build the purine ring, producing glutamate that Complex II must oxidize to sustain purine synthesis. This connection translated to a metabolic vulnerability whereby increasing intracellular glutamate levels suppresses purine production and sensitizes AML to Complex II inhibition. In mouse models, targeting Complex II triggered rapid disease regression and extended survival in aggressive AML. The clinical relevance of this pathway emerged in human studies, where higher Complex II gene expression correlates with both resistance to mitochondria-targeted therapies and worse survival outcomes specifically in AML patients. These findings establish Complex II as a central regulator of *de novo* purine biosynthesis and identify it as a promising therapeutic target in AML.

## Introduction

Cells orchestrate their functions through intricate networks of interacting genes. Co-essentiality mapping reveals these networks by identifying genes that share similar patterns of essentiality across conditions^4–6^. While this approach has successfully uncovered important gene pairs^6,7^, scaling to genome-wide analysis for human genomic research presents nearly 400 million possible interactions. In contrast, mapping connections between gene pathways may provide an opportunity to organize co-essentiality data into higher levels to identify associations less apparent with single gene-gene queries. Interrogating pathway-level interactions may be of particular interest in metabolism research, as a growing number of studies demonstrate that metabolic pathways connect to and regulate each other in unexpected ways. Specifically, mitochondrial electron transport chain (ETC) complexes have been shown to have critical roles in cancer by supporting amino acid^8–10^, nucleotide^11–13^, and epigenetic^14,15^ processes. Here, we developed a pathway co-essentiality mapping tool to determine pathway-pathway interactions across the genome, focusing particularly on ETC components. This approach revealed an unexpected discovery: Complex II (succinate dehydrogenase, SDH) directly regulates purine synthesis in AML, driving disease progression through this previously unknown metabolic link.

### Complex II has unique pathway associations

To map pathway-level connections, we analyzed nearly 3,000 gene sets from multiple annotation systems: Canonical pathways, Gene Ontology (GO) pathways, Transcription Factor targets, and Cell Type signatures. These sets encompassed approximately 15,000 expressed genes^16^. To evaluate pathway-level connections, we developed a systematic approach using CRISPR-based dependency scores from the Cancer Dependency Map^3^. First, we extracted the top Principal Components (PCs) for each pathway. Then, we used Canonical Correlation Analysis (CCA)^17^ to find maximum correlations between pathway PCs (Fig. 1a). This revealed distinct patterns of pathway co-essentiality across major biological domains – Biological Process, Molecular Function, and Cellular Components (Extended data Fig. 1a-c). Importantly, embedding genes as pathways yielded stronger co-essentiality correlations than individual gene-pathway comparisons (Extended Data Fig 1d-e).

**Fig. 1:**
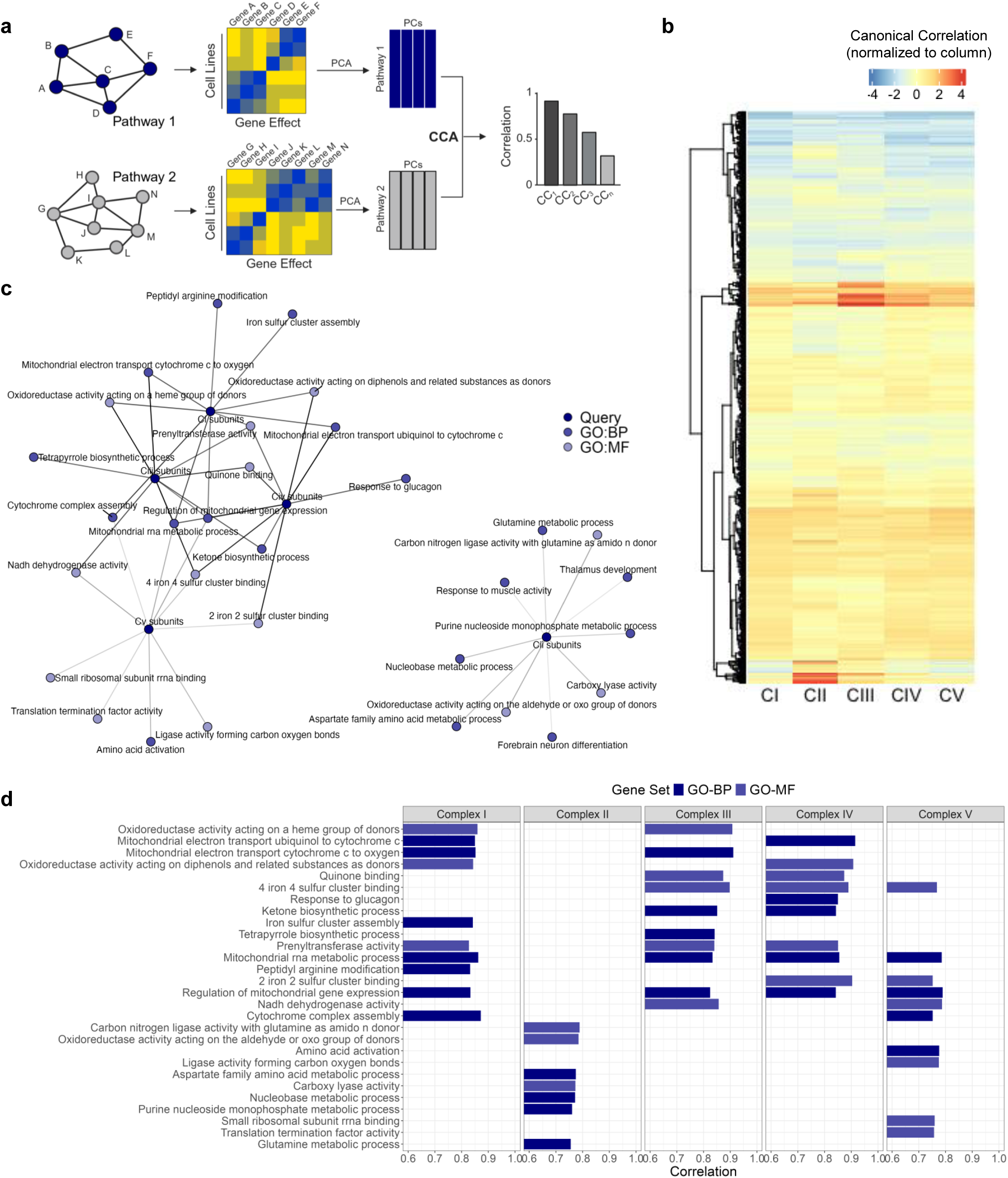
Pathway coessentiality mapping reveals a unique association between Complex II and nucleotides. a) Development of pathway mapping model. Gene sets representing ∼15,000 genes were used to select essentiality data from the Cancer Dependency Map. For each Gene Ontology (GO) pathway, the first four Principal Components (PCs) were extracted and Canonical Correlation Analysis (CCA) was used to determine correlations between pathways. b) Heatmap of the canonical correlation matrix for each ETC complex across 2,827 pathway gene sets. K-nearest neighbors hierarchical clustering was used to group pathways by their co-essentiality patterns (side). The color of each cell represents the relative canonical correlation strength for each pathway-complex comparison. c) Network diagram showing the connectedness of ETC complexes with GO pathways, with each ETC complex or pathway represented as a node, and each connection as an edge. Notably, a larger sub-network includes Complexes I, III, IV, and V and related pathways, whereas a smaller sub-network includes only Complex II and its unique pathways. d) Top pathway associations for each ETC complex, with shared pathways shown for Complexes I, III, IV and V and distinct pathways for Complex II. For panels c-d, color of nodes or bars represents GO pathway type (Biologic Process: BP or Molecular Function: MF) as indicated.

The ETC served as an ideal test case for this approach. Its five protein complexes coordinate hydrogen pumping with ATP production during oxidative phosphorylation (OxPhos) and support various biosynthetic functions^8,9^. We next analyzed how each ETC complex (I-V) connected to cellular pathways by examining their subunits collectively (Extended Data Fig 1f). Hierarchical clustering revealed Complex II stood apart from the other ETC components (Fig. 1b). While Complexes III and IV showed strong similarity in their pathway connections, as did Complexes I, III, IV, and V - matching their known ability to form supercomplexes^18^ - Complex II showed a unique pattern. This distinction appeared both in pathway associations and in direct ETC gene co-essentiality analysis (Extended Data Fig. 1g). Network visualization reinforced this finding, showing that Complexes I, III, IV and V formed a tight cluster through shared co-essential pathways, while Complex II and its associated pathways formed a separate, distinct network (Fig. 1c).

Complex II oxidizes succinate while reducing ubiquinone^19^. Our computational analysis revealed 31 cellular pathways with unique Complex II connectivity (Extended Data Table 1). The strongest connections emerged with nucleotide and amino acid metabolism - specifically the “Nucleobase Metabolic Process,” “Purine Nucleobase Monophosphate Metabolic Process,” “Aspartate Family Metabolic Process,” and “Glutamine Metabolic Process.” No other ETC complex shared these metabolic connections (Fig. 1d). This distinct pathway signature pointed to a previously unrecognized role for Complex II in cancer cell metabolism via the potential regulation of nucleotide biosynthesis.

### Complex II regulates purines to maintain AML survival

We analyzed cancer dependency data from DepMap to understand which cancers rely most on ETC complexes, which revealed that blood cancers have greater sensitivity to Complex II disruption (Extended Data Fig. 2a). Specifically, both myeloid and lymphoid cancers depend strongly on Complex II for survival. Further examination of specific cancer subtypes highlighted Complex II’s critical role in multiple types of lymphomas and leukemias, including AML and related disease such as Myelodyplastic Syndromes and Myeloproliferative Neoplasms (Extended Data Fig. 2b).

To validate this computational finding, we screened a panel of 72 diverse cancer cell lines for sensitivity to Complex II inhibition. We used 2-thenoyltrifluoroacetone (TTFA), a potent Complex II inhibitor, at a dose (200 μM) that effectively blocks Complex II activity^20^. Consistent with our computational analyses, cell lines derived from the hematologic malignancies, including AML and diffuse large B cell lymphoma (DLBCL), were most sensitive to Complex II inhibition (Fig. 2a). We also confirmed the sensitivity profile of a subset of cancer cell lines with an additional inhibitor, 3-nitroproprionic acid (3-NP), to ensure that results were representative of on-target Complex II inhibition (Extended Data Fig. 2c). We found that Complex II dependency was not driven by differences in target inhibition, as measured by succinate accumulation or by ATP loss (Extended Data Fig 2d-e). These results confirmed our computational prediction that blood cancers, including AML, have a specific vulnerability to Complex II inhibition that distinguishes them from other cancer types.

**Fig. 2:**
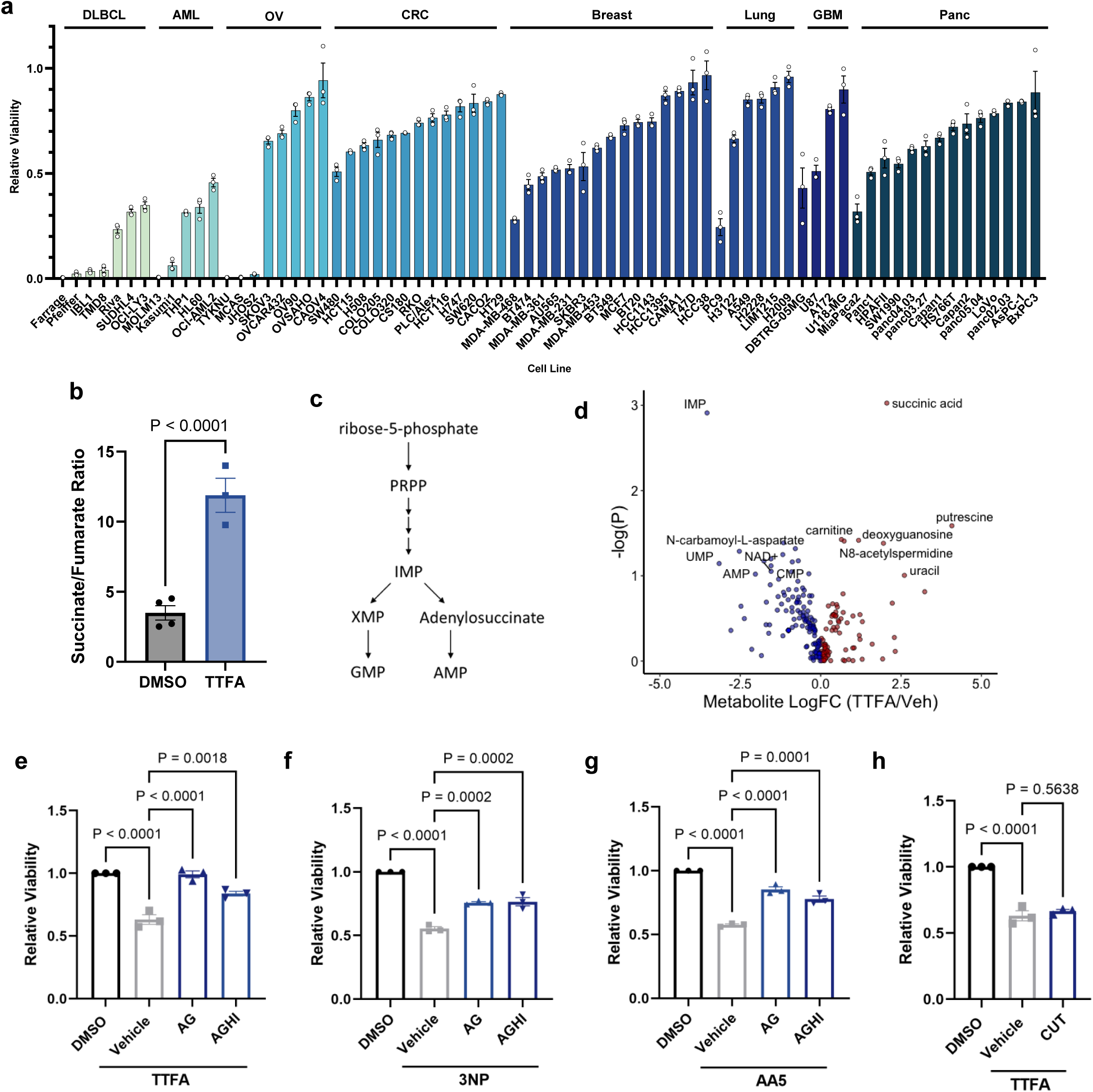
Complex II regulates purine levels to maintain AML survival. a) Relative viability for 72 cancer cell lines after 3 days of Complex II inhibition with TTFA. Cell lines are grouped by cancer cell lineage according to CCLE annotations (DLBCL, Diffuse Large B-cell Lymphoma; OV, Ovarian; CRC, Colorectal Carcinoma; GBM, Glioblastoma Multifome; Panc, Pancreatic). b) Metabolite ratio of succinate to fumarate after 24hr in control conditions (n=4) or with Complex II inhibition (n=3). c) Diagram overview of *de novo* purine biosynthesis. The purine ring is synthesized on a PRPP backbone that is derived from glucose through ribose-5-phosphate. d) Volcano plot showing the effect of Complex II inhibition on steady state metabolite levels in MOLM-13 cells. e-g) Effect of exogenous purine add-back of adenine (A), guanine (G), hypoxanthine (H) or inosine (I) (each 30 μM) on AML viability following Complex II inhibition with (e) TTFA, (f) 3NP, or (g) AA5 in OCI-AML2 cells. h) Effect of pyrimidine addback on MOLM-13 AML cell viability with Complex II inhibition. For all plots, data are shown as mean ± s.e.m. of 3 independent experiments in. *P* values were determined by unpaired, two-tailed Student’s t-test (b) or by one-way ANOVA with Tukey’s post-hoc test (e-h).

To explore the metabolic pathways that emerged as uniquely co-essential with Complex II, we performed LC-MS metabolic profiling. Complex II inhibition increased the succinate:fumarate ratio (Fig. 2b) and reduced levels of multiple nucleotides, particularly *de novo* purines like inosine monophosphate (IMP) (Fig. 2c-d and Extended Data Fig. 3a-c). This suggested that AML’s sensitivity to Complex II inhibition stems from purine depletion. Cells can naturally compensate for reduced purine synthesis through the salvage pathway, which converts exogenous purines - adenine (A), guanine (G), hypoxanthine (H), and inosine (I) - into their respective nucleotides. Previous studies showed that purine salvage can rescue cells from purine synthesis inhibitors^21,22^. Further, Complex I inhibition activates this salvage pathway through HPRT1^11^. Thus, we tested if supplementation with purines (AG or AGHI) rescued cells from Complex II inhibition by TTFA. First, we established that purines rescued cell growth from treatment with the purine synthesis inhibitor lometrexol (LTX) (Extended Data Fig. 3d). As an additional control, pyrimidines rescued cells from the DHODH inhibitor brequinar but not from purine synthesis inhibition (Extended Data Fig. 3d-e). Remarkably, external purines also rescued AML cells from Complex II inhibition by TTFA (Fig. 2e). This rescue extended to two structurally distinct Complex II inhibitors, atpenin A5 (AA5) and 3-NP (Fig. 2f-g). However, pyrimidines failed to rescue cells from TTFA treatment despite their effectiveness against pyrimidine synthesis inhibition (Fig. 2h). These results establish that Complex II maintains AML survival specifically through purine regulation.

### Complex II supports *de novo* purine synthesis

Next, we used stable-isotope tracing with labeled glucose or glutamine to uncover how Complex II influences purine biology (Fig. 3a). Complex II inhibition caused striking metabolic changes. First, ^13^C_6_-glucose-derived carbon accumulated in succinate and upstream TCA cycle metabolites (α-ketoglutarate and citrate), indicating blocked oxidative glucose metabolism (Fig. 3b-c and Extended Data Fig. 4a-d). This blockade forced glucose-derived carbon into alternative pathways, increasing glycolytic metabolites pyruvate and lactate (Extended Data Fig. 4e-f).

**Fig. 3:**
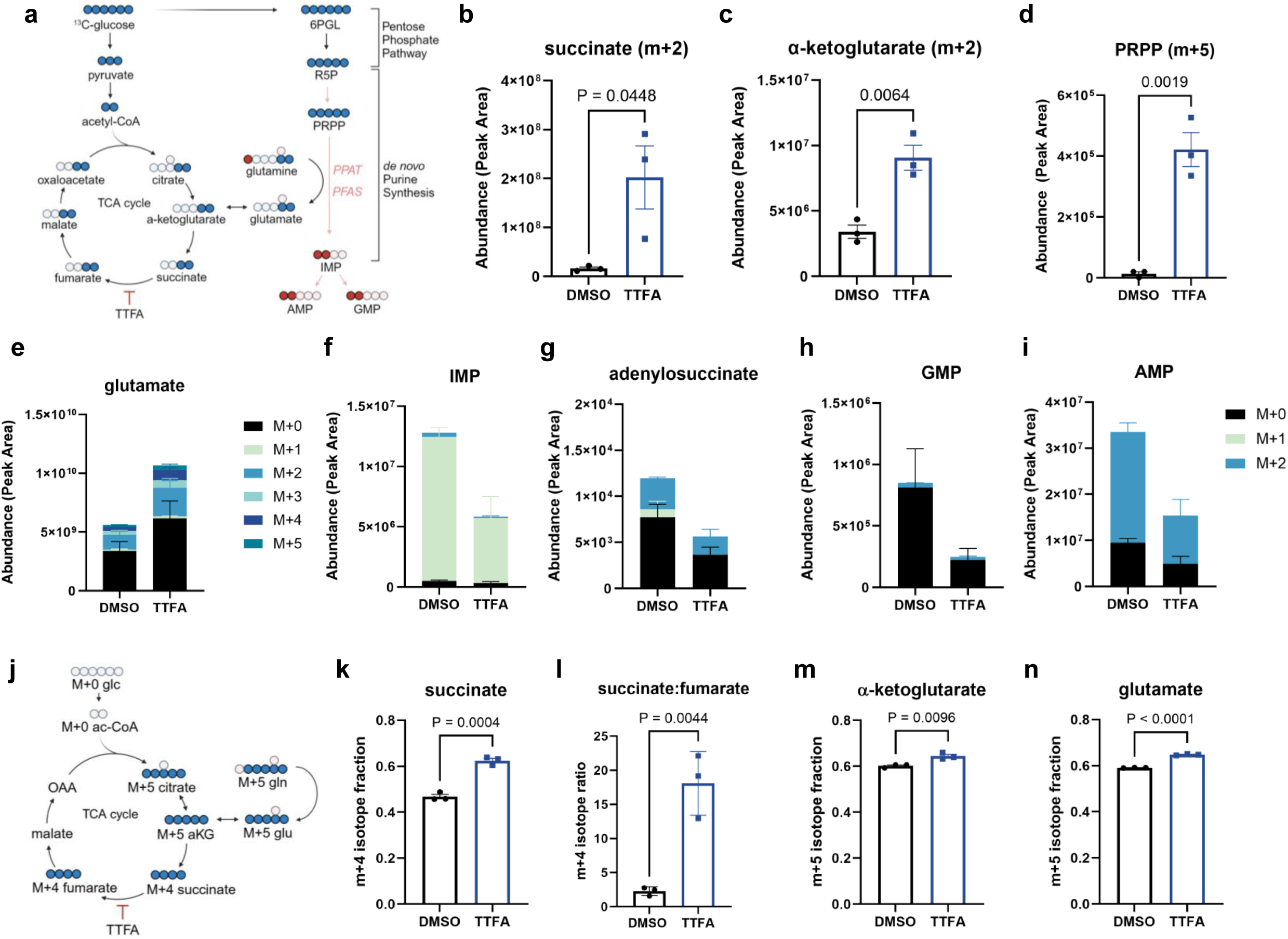
Complex II supports *de novo* purine biosynthesis. a) Overview of metabolic pathways interrogated with stable isotope tracing studies using ^13^C_6_-glucose or ^15^N_1_-glutamine. Relevant metabolites for ^13^C_6_-glucose tracing are shown in blue, and ^15^N_1_-glutamine tracing are shown in red. b-c) Effect of Complex II inhibition on ^13^C_6_-glucose derived carbon in (b) m+2 succinate, (c) m+2 α-ketoglutarate. Results show mean ± s.e.m. for all isotope peak areas (n=3 replicates per group). d) Effect of Complex II inhibition on ^13^C_6_-glucose derived m+5 PRPP. e) Effect of Complex II inhibition on ^13^C_6_-glucose derived glutamate. Results show mean ± s.e.m. for all isotope peak areas, n=3 replicates per group. f-i) Effect of Complex II inhibition on ^15^N_1_-glutamine nitrogen into *de novo* purines including (f) IMP, (g) adenylosuccinate, (h) GMP and (i) AMP in MOLM-13 cells. Data show mean ± s.e.m. for n=3 replicate experiments per group. j) Overview of possible routes of carbon distribution in oxidative or reductive TCA cycle reactions following anapleurosis of ^13^C_5_-glutamine. k-n) Effect of Complex II inhibition on ^13^C_5_-glutamine carbon for the following metabolite isotopes: (k) m+4 succinate, (l) m+4 succinate:m+4 fumarate ratio, (m) m+5 α-ketoglutarate and (n) m+5 glutamate. Bars show fractional abundance for indicated metabolite and isotope. P values are shown for unpaired t-tests.

Glucose enters purine synthesis through the Pentose Phosphate Pathway. This pathway produces phosphoribosyl pyrophosphate (PRPP), which forms the sugar backbone for all nucleotides. Given our observation that TTFA treatment reduces purine levels (Fig. 2d), we examined PRPP metabolism. Both Complex II inhibition (TTFA) and direct purine synthesis inhibition (LTX) caused PRPP accumulation (Fig. 3d and Extended Data Fig. 4g). Complex II inhibition also increased glucose-derived carbon in other purine building blocks - glycine, serine, and aspartate (Extended Data Fig. 4h-k). These results demonstrate that blocking Complex II causes purine precursors to accumulate while final purine products decline.

Previous work showed that in osteosarcoma cells, aspartate or the electron acceptor α- ketobutyrate (AKB) can rescue Complex II inhibition by AA5^23^. However, in AML cells, we observed a different pattern. Aspartate synthesis remained intact during Complex II inhibition, as both oxidatively-derived aspartate (m+2) and pyruvate carboxylase-derived aspartate (m+3) levels stayed constant with TTFA treatment (Extended Data Fig. 4l-m). Unlike Complex I inhibition, which aspartate can rescue, neither aspartate nor AKB rescued AML cells from Complex II inhibition (Extended Data Fig. 4n-p). This ruled out aspartate deficiency as the cause of purine suppression in AML.

Glutamine emerged as a key player in this metabolic circuit. During purine synthesis, glutamine provides nitrogen to build the purine ring, which contributes 50% of the nitrogen atoms in the final purine structure ^24^. This occurs at two critical steps: first when PRPP amidotransferase (PPAT) converts PRPP to phosphoribosyl-β amine (PRA), and again when FGAM synthetase converts formylglycinamide ribonucleotide (FGAR) to formylglycinamidine ribonucleotide (FGAM) (Fig. 3a and Extended Data Fig. 4h). Each time a nitrogen is transferred, glutamine is converted to glutamate^25^.

Our ^13^C_6_-glucose tracing revealed Complex II inhibition caused glutamate accumulation while glutamine levels remained stable (Fig. 3e and Extended Data Fig. 5a-b). This aligned with our earlier finding of a computational link between Complex II and glutamine nitrogen donation (Fig. 1c-d). Using ^15^N_1_-glutamine-amide tracing, we confirmed that Complex II inhibition specifically impaired glutamine nitrogen donation to purines, shown by decreased m+1 and m+2 isotopes of IMP, AMP, GMP, and adenylosuccinate (Fig. 3f-i). This matched the effect of direct purine synthesis inhibition with LTX (Extended Data Fig. 5c-f). Importantly, pyrimidine synthesis remained largely unaffected (Extended Data Fig. 5g-m), consistent with the inability of exogenous pyrimidines to rescue Complex II inhibition in AML (Fig. 2h).

While some ETC mutations increase glutamine anaplerosis into the TCA cycle^26^, Complex II inhibition had the opposite effect. Using ^13^C_5_-glutamine tracing (Fig 3j), we found that TTFA treatment blocked glutamine carbon oxidation in the TCA cycle. This caused accumulation of m+4 succinate, m+5 α-ketoglutarate and m+5 glutamate, plus an increased m+4 succinate to m+4 fumarate ratio (Fig. 3k-n). Complex II inhibition also reduced glutamine’s reductive carboxylation, shown by decreased m+5 citrate (Extended Data Fig. 5n). In sum, these data demonstrate that Complex II inhibition creates a metabolic bottleneck by blocking carbon flow through the TCA cycle while simultaneously disrupting glutamine’s critical nitrogen donation to purine synthesis.

### Glutamate links Complex II to purines

To understand the broader metabolic impact of Complex II inhibition, we analyzed steady-state metabolites after TTFA treatment (Extended Data Fig. 6a and Supplementary Table 2). This analysis revealed widespread effects beyond nucleotide metabolism, particularly in amino acid pathways involving alanine, aspartate, and glutamate. We also observed changes in arginine and proline metabolism, aligning with recent findings that OxPhos activity reciprocally regulates proline and ornithine synthesis^27^.

Glutamate emerged as a central player in these metabolic changes. Beyond providing carbon to the TCA cycle, glutamate serves as a substrate for glutathione synthesis and regulates cystine import through system Xc^28^ (Fig. 4a). Complex II inhibition significantly altered glutathione metabolism in AML cells, causing them to redirect both glucose and glutamine carbon into glutathione synthesis (Extended Data Fig. 6b-c).

**Fig. 4:**
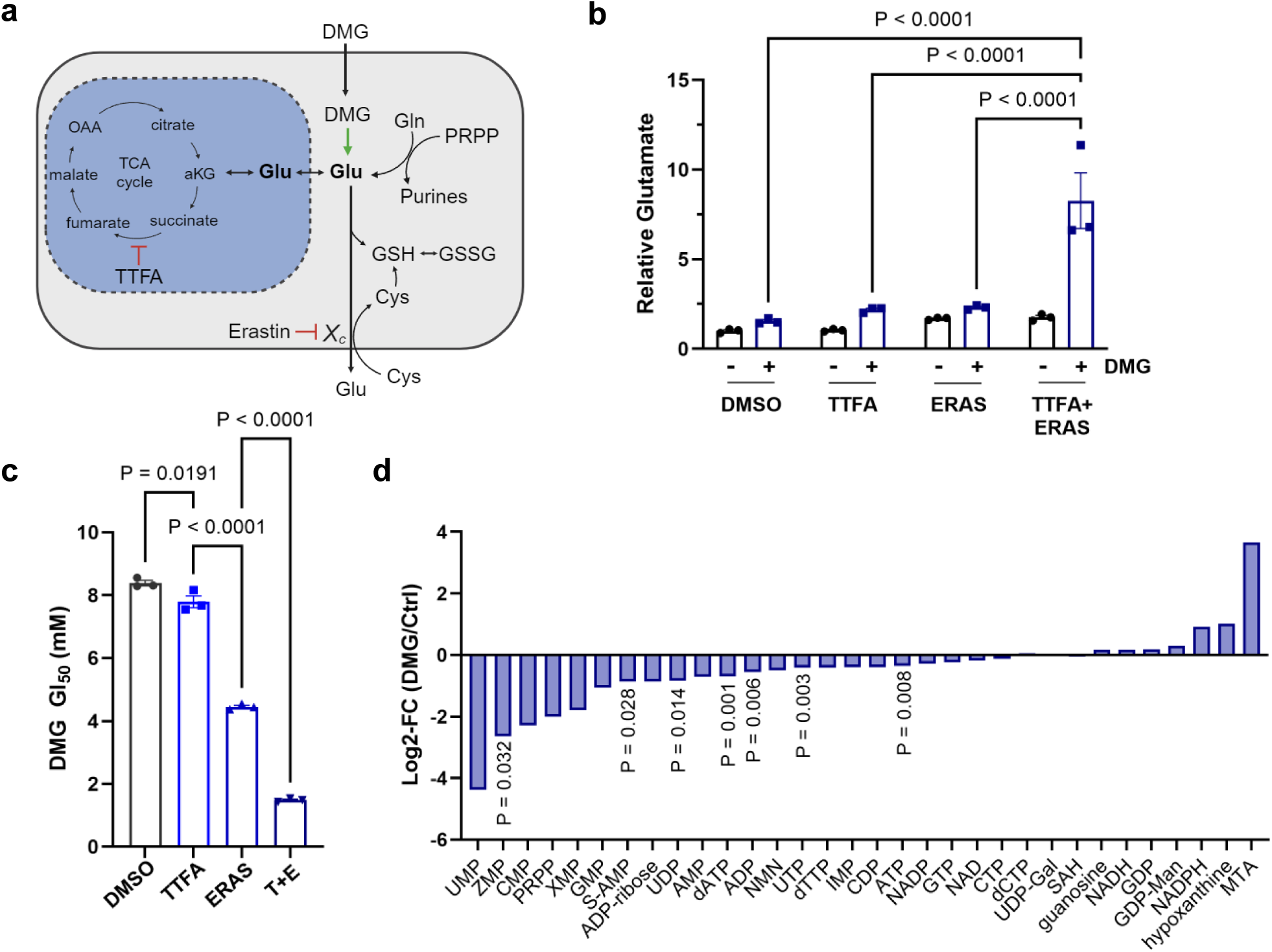
Glutamate potentiates Complex II inhibition and reduces *de novo* purines. a) Diagram showing potential metabolic fates of glutamate as related to Complex II inhibition, highlighting that glutamate produced from *de novo* purine biosynthesis can be used for TCA cycle anapleurosis. Alternatively, glutamate can be exported through system Xc (which is targeted by erastin) or used for glutathione biosynthesis that is coupled to cystine import. Abbreviations: DMG, dimethyl-glutamate; GSH, reduced glutathione; GSSH, glutathione disulfide; Gln, glutamine; Glu, glutamate; Cys, cystine; aKG, alpha-ketoglutarate; OAA, oxaloacetate). b) Glutamate loading with DMG in combination with Complex II inhibition (TTFA, T) or system Xc inhibition by erastin (ERAS, E) increases intracellular glutamate concentration. Data show mean ± s.e.m. from n=3 replicates and were analyzed using one-way ANOVA with a Tukey post-hoc test. c) Effect of Complex II inhibition alone or in combination with erastin on the 50% growth inhibitory dose of DMG in MOLM-13 cells. d) Waterfall plot showing the effect of DMG treatment on nucleotide metabolites in MOLM-13 AML cells. Data show log-2 fold change for n=3 replicates and were analyzed with ANOVA and a post-hoc Tukey test.

These findings pointed to a cellular adaptation where AML cells must manage excess glutamate during Complex II inhibition to prevent it from interfering with purine synthesis. We confirmed elevated intracellular glutamate across multiple AML cell lines after TTFA treatment (Extended Data Fig. 6d-f). To further investigate this glutamate-purine connection, we used three approaches to increase cellular glutamate: TTFA (Complex II inhibition), erastin (system Xc inhibition), and dimethyl-glutamate (DMG, a cell-permeable glutamate). Both system Xc inhibition and DMG worked synergistically with TTFA to elevate cellular glutamate levels (Fig. 4b). Next, we reasoned if glutamate accumulation drives purine synthesis suppression during Complex II inhibition, then increasing glutamate should enhance TTFA’s effects. Indeed, erastin treatment increased cellular sensitivity to DMG and TTFA, shown by a lower GI_50_ (Fig. 4c and Extended Data Fig 6g-i). Finally, direct metabolomic analysis confirmed that DMG treatment alone reduced intracellular purine levels (Fig. 4d).

These results reveal a metabolic vulnerability in AML where Complex II inhibition triggers glutamate accumulation, which suppresses purine synthesis and enhances cell death. This mechanism suggests new therapeutic strategies combining Complex II inhibition with approaches that increase cellular glutamate.

### Complex II is essential for AML progression

Our metabolic studies revealed that Complex II supports AML survival through purine biosynthesis regulation. In particular, Complex II function requires its catalytic subunit *SDHB*, which handles Fe-S mediated electron transfer to ubiquinone. To test whether this mechanism is observed in physiological mouse models, we examined Complex II inhibition in a syngeneic *MLL-AF9*-driven mouse model of AML (Fig. 5a). This model replicates human AML driven by the t(9;11)(p22;q23) translocation in blood stem cells^29^. We depleted *Sdhb* through doxycycline-induced shRNA, which potently reduced both *Sdhb* expression. Remarkably, lower SDHB reduced AML burden in mouse bone marrow and spleen and significantly extended mouse survival (Fig. 5b-d and Extended Data Fig. 7a). These data demonstrate SDHB influences AML survival *in vitro* and *in vivo*.

**Fig. 5:**
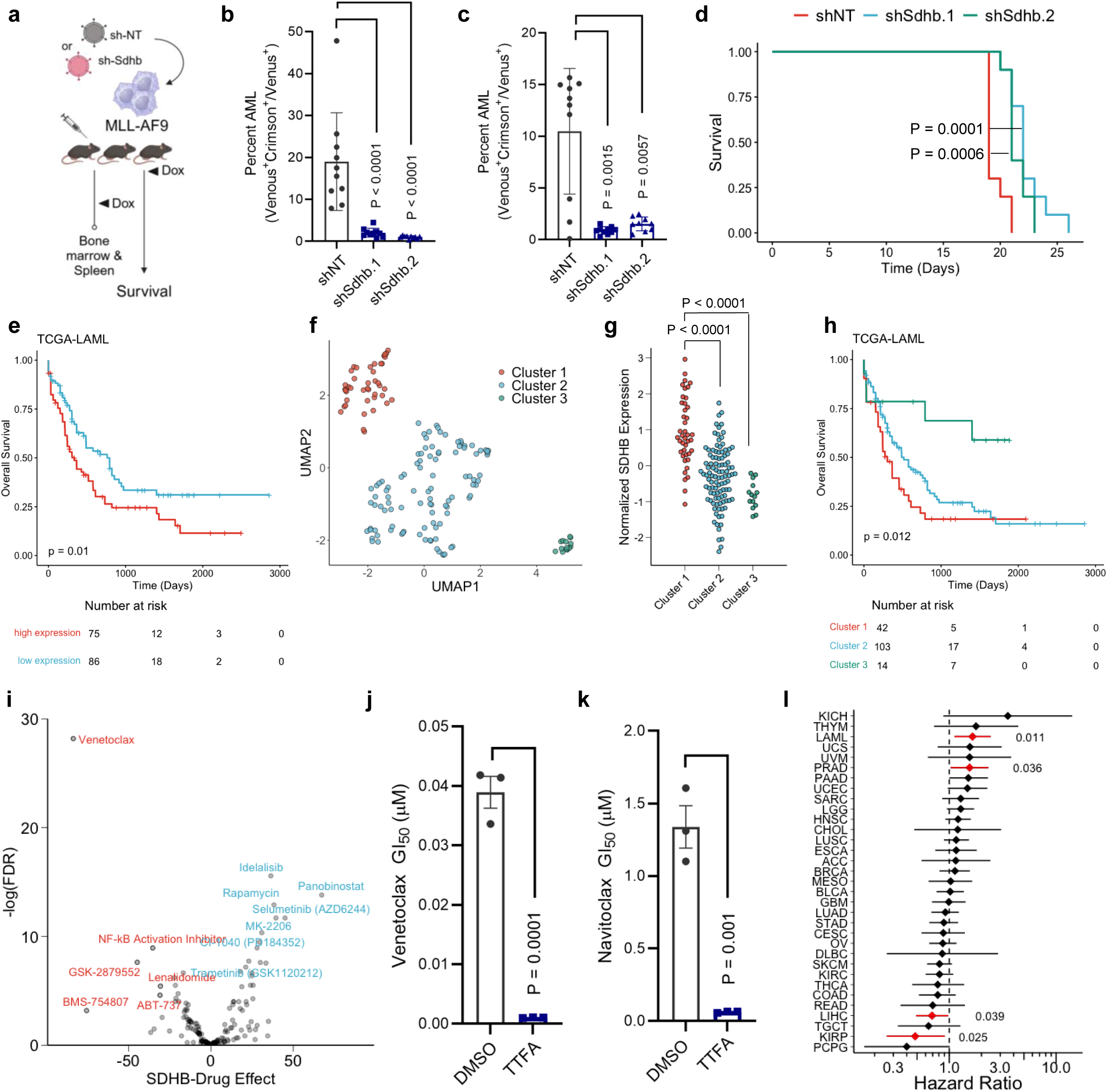
SDHB is a high risk molecular feature that is essential for AML *in vivo*. a) Experimental model for SDHB knockdown in the MLL-AF9 mouse. Cells were transfected with retroviral vectors expressing doxycycline-inducible non-targeting (shNT) or Sdhb-targeting (shSdhb) small hairpin RNA and transplanted into recipient mice for bone marrow, spleen, or survival analysis. b) Bone marrow AML burden after doxycycline treatment in the MLL-AF9 model. AML actively expressing MLL-AF9 and shSdhb or shNT (Venus^+^Crimson^+^) were compared to all cells expressing MLL-AF9 (Venus^+^). c) Spleen AML burden after doxycycline treatment. d) Survival of MLL-AF9 leukemic mice after shNT or shSdhb induction with doxycycline (n=10 mice per treatment group for each experiment in panels b-d). e) Kaplan Meier curve showing overall survival of the TCGA AML cohort, stratified by *SDHB* expression above or below the mean. f) UMAP analysis of TCGA-AML data, showing three clusters based on whole transcriptome data. g) *SDHB* expression within the 3 TCGA UMAP clusters from (f). h) Overall survival for AML UMAP clusters, with cluster 1 having the highest *SDHB* expression and the poorest overall survival. i) Volcano plot showing the association between *SDHB* gene expression and inhibitor sensitivity in the Beat AML cohort^33^. Higher *SDHB* expression in patient blasts was associated with resistance to inhibitors on the left of the plot and sensitivity to inhibitors on the right. j-k) Effect of Complex II inhibition (200 μM TTFA) on the GI_50_ of MOLM-13 AML cells to the BCL-2 inhibitors (j) venetoclax and (k) navitoclax. l) Hazard Ratio plot showing the risk of death associated with elevated *SDHB* expression across all TCGA cancer types. Red indicates cancer types with significant hazard ratios, with p value shown at the side.

These findings prompted us to investigate Complex II’s role in human AML. While many cancers depend on OxPhos for growth and metastasis^30^, mitochondrial therapy targeting BCL-2 has shown particular promise in leukemia^31,32^. Survival analysis of The Cancer Genome Atlas (TCGA) data revealed that high expression of Complex II subunits *SDHB* or *SDHA* was associated with poor AML patient survival (Fig. 5e and Extended Data Fig. 7b, respectively). Extending this approach, we performed a dimension reduction analysis on TCGA-LAML transcriptomes using UMAP and found three distinct patient clusters (Fig. 5f), again with the poorest-surviving group showing the highest *SDHB* expression (Fig. 5g-h).

Next, we turned to the Beat AML dataset^33^ to determine if *SDHB* expression was also associated with treatment response. We performed a regression analysis to identify associations between drug sensitivity and *SDHB* expression. We found high *SDHB* expression correlated with resistance to 19 targeted molecules, most notably the BCL-2 inhibitor venetoclax (Fig. 5i and Extended Data Fig. 7c-d). This suggested a therapeutic opportunity by combining Complex II and BCL-2 inhibition. Indeed, TTFA treatment sensitized AML cells to both venetoclax and navitoclax (Fig. 5j-k and Extended Data Fig. 7e-h).

Complex II’s role in AML stands out among human cancers. Calculating the Hazard Ratio (HR) of high *SDHB* expression across 33 cancer types in TCGA, we found high *SDHB* expression significantly increased mortality risk in only two cancers, with AML being one (Fig. 5l). This contrasts sharply with tumors like pheochromocytoma, paraganglioma (PCPG), and renal papillary carcinoma (KIRP), where high *SDHB* expression was associated with improved survival. In these cancers, Complex II acts as a tumor suppressor, and its loss drives hereditary pheochromocytoma paraganglioma syndromes^34,35^. This opposite effect, where Complex II promotes disease in AML but suppresses it in other cancers, highlights its unique role in AML progression and suggests targeted therapeutic opportunities.

## Discussion

We developed a new algorithm that moves beyond established gene-gene co-essentiality mapping and reveals pathway-pathway interactions across biological pathways. We applied this new approach to examine relationships among the five complexes of the ETC, and discovered a novel association between Complex II and purine synthesis. Furthermore, we demonstrate that Complex II directly regulates purine synthesis to maintain cell proliferation. Mechanistically, we provide evidence that an important function of Complex II is to oxidize the carbon from glutamine. Upon Complex II inhibition, glutamine carbon (i.e. glutamate) oxidation is reduced, which causes negative feedback and a block of purine synthesis.

Our finding that Complex II regulates purine synthesis adds to a growing body of work linking the ETC and TCA cycle to non-canonical functions, particularly nucleotide synthesis. Landmark studies revealed that the major role of mitochondrial Complex I in maintaining proliferation is not through ATP production but rather through NAD^+^/NADH-dependent aspartate biosynthesis^36,37^. The reduction in NAD^+^/NADH also results in TCA cycle inhibition and depletion of asparagine-mediated nucleotide production^38^. Recently, a number of studies have connected the ETC and TCA cycle enzymes directly to purine synthesis, including complex I^11^, OGDH^39^, and fumarate hydratase (FH)^40^. Models of FH deficiency were shown to block purine synthesis via fumarate accumulation driving the reversal of adenylosuccinate lyase (ADSL)^40^. This observation supports our proposed model of glutamate accumulation caused by Complex II inhibition blocking purine synthesis, suggesting a novel mechanism of cellular sensing. Complex II is unique among the ETC complexes since it has a dual role in the TCA cycle. As the TCA cycle is a known biosynthetic metabolic hub, it is noteworthy that FH and SDH inhibition both purportedly suppress purine pathway enzymes through substrate accumulation, as opposed to mechanisms related to alterations in electron acceptors or aspartate/asparagine synthesis.

Our work also highlights the potential for Complex II to be used therapeutically against cancer. As previously mentioned, targeting OXPHOS for cancer therapy has proven to be challenging, mainly due to a lack of therapeutic window and dose-limiting toxicities that are unable to achieve clinical benefit. In particular, Complex I inhibition has recently been associated with neurotoxicities^32^. However, a few lines of evidence suggest that Complex II may be a viable therapeutic target.

First, we find a subset of cancer cell lines are exquisitely dependent upon Complex II, suggesting there may be an exploitable therapeutic window in specific cancers. Our data suggest this may be the case for AML in particular, where OxPhos inhibition could be combined with by venetoclax and azacitidine – agents well known to selectively eradicate leukemic stem cells^41^. We show that Complex II inhibition sensitizes AML cells to BCL-2 inhibition, providing a possible area of exploration in venetoclax-resistant disease^42^. In addition to AML, our data also point to a broader essential role for Complex II in hematolymphoid malignancies, including B-cell acute lymphoblastic leukemia and certain lymphomas, particularly anaplastic large cell lymphoma (ALCL) and DLBCL. These findings highlight the need for further studies on whether the essential functions of Complex II are shared amongst subsets of cancers. In fact, using TCGA data, we show that the increased risk of mortality with elevated *SDHB* expression is not a pan-cancer phenomenon, further implicating Complex II in AML and distinguishing it from tumors in which Complex II acts as a tumor suppressor.

Second, several studies have shown that Complex II ablation (*SdhD*) is tolerable in adult mice^43,44^. Models of partial or heterozygous whole-body knockout of *SdhD* and *SdhC* in adult mice reduced SDH activity and were well-tolerated, with no major pathophysiology observed other than reduced carotid body volume ^43,44^. Of note, germline mutations in *SDH* subunits predispose humans to a small subset of cancers, including pheochromocytoma, paraganglioma, gastrointestinal stromal, and renal tumors, which are manifested upon loss of the wild-type allele^45^. However, *Sdh* loss in mice was not associated with tumor development^43,44^. Using TCGA data, we have found that the risk of elevated *SDHB* expression is cancer-dependent, suggesting that Complex II targeting may be more tissue selective and may broaden the therapeutic window. Additionally, mitochondrial-targeted lonidamine (LND), a clinically safe glycolysis inhibitor found to inhibit Complex II, was well-tolerated in mice and inhibited the growth of lung cancer xenografts ^46^. However, other pharmacological inhibitors of Complex II, 3-NP and malonate, have been investigated *in vivo* and found to cause striatal defects analogous to those in Huntington’s disease with long-term use ^47,48^. Since neurotoxicity is a primary concern with ETC inhibition, this raises the possibility of creating a brain-impermeant Complex II inhibitor to retain its anti-tumorigenic properties while avoiding serious side effects. Finally, we note that OGDH ablation was reported to have similar inhibitory effects on purine synthesis, but ascribed to different mechanisms^49^. Because knockdown of *Ogdh* in mice results in mild side effects but is generally well-tolerated^49^, it may be an alternative therapeutically targetable node in Complex II-dependent cancers. Overall these data support a key role of the TCA cycle and ETC in supporting purine biosynthesis in AML and other hematolymphoid malignancies.

## EXTENDED DATA LEGENDS

**Extended Data 1 (Related to Fig. 1).**
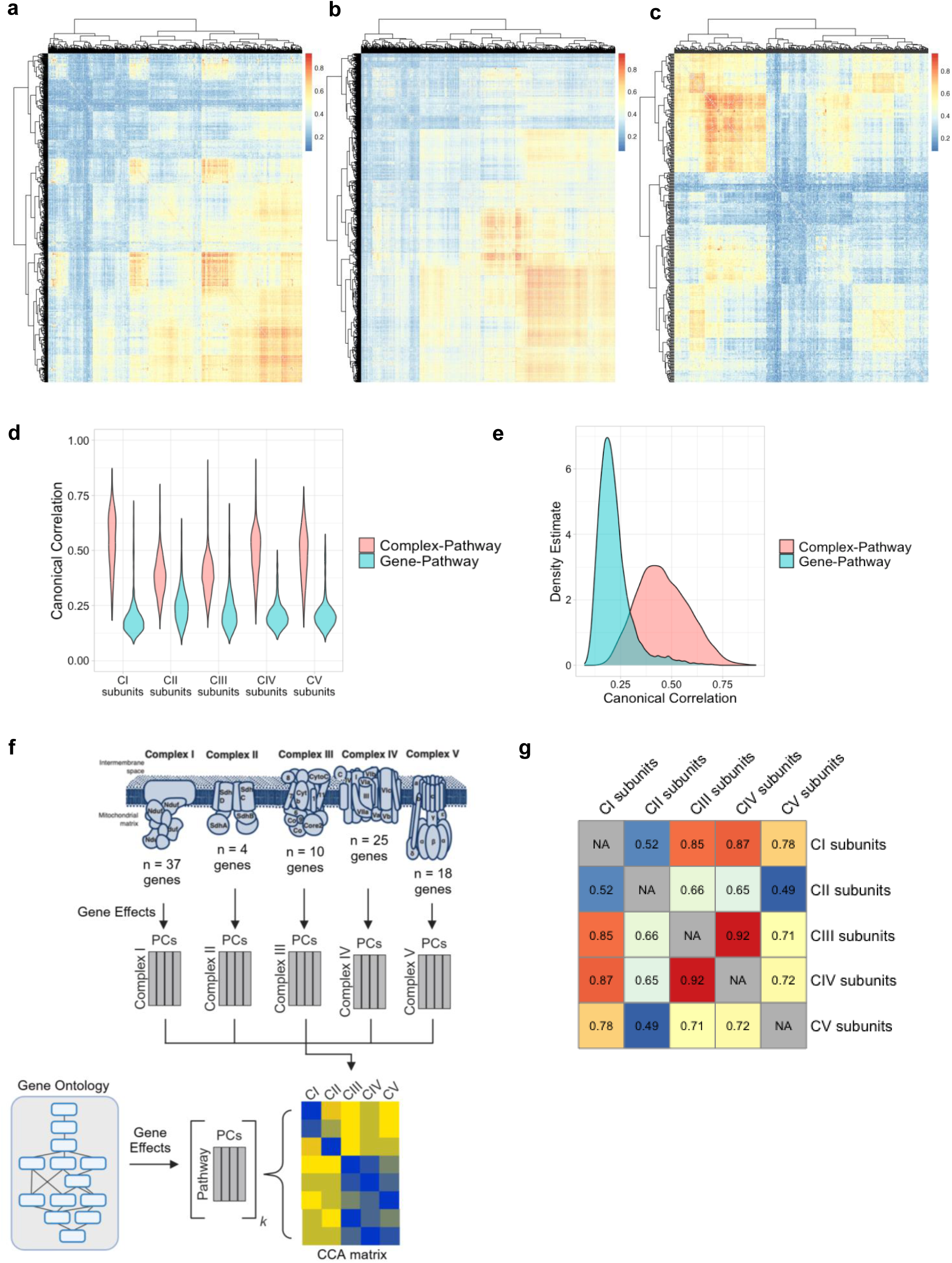
a-c) Heatmaps depicting pathway-pathway correlation matrices for Gene Ontology (GO) (a) Molecular Function (MF), (b) Biologic Process (BP), and (c) Cellular Component (CC) categories. d) Violin plot showing the effect of Canonical Correlation Analysis (CCA) on the correlation strengths between each ETC complex and all pathways. Pink plots represent the correlation values when comparing all pathways to ETC complexes using principal components (Canonical Correlation), teal plots represent the distribution of correlations when comparing individual genes making up each ETC Complex without first extracting principal components. Correlation strength was consistently stronger when PCs were first extracted to represent ETC genes *en bloc*. e) Overall correlation distribution between pathway-pathway and gene-pathway comparisons for all ETC-pathway correlations from (d). f) Overview of the analysis to determine canonical correlation between ETC complexes and GO pathways showing the number of genes used to represent each ETC complex as a pathway, the representation of ETC complexes or GO pathways in PCs, and the CCA-generated matrix of complex-pathway correlations. g) Correlation analysis between ETC complexes, with the color of each cell and associated correlation score (shown within each cell) representing the strength of complex-complex associations.

**Extended Data 2 (Related to Fig. 2).**
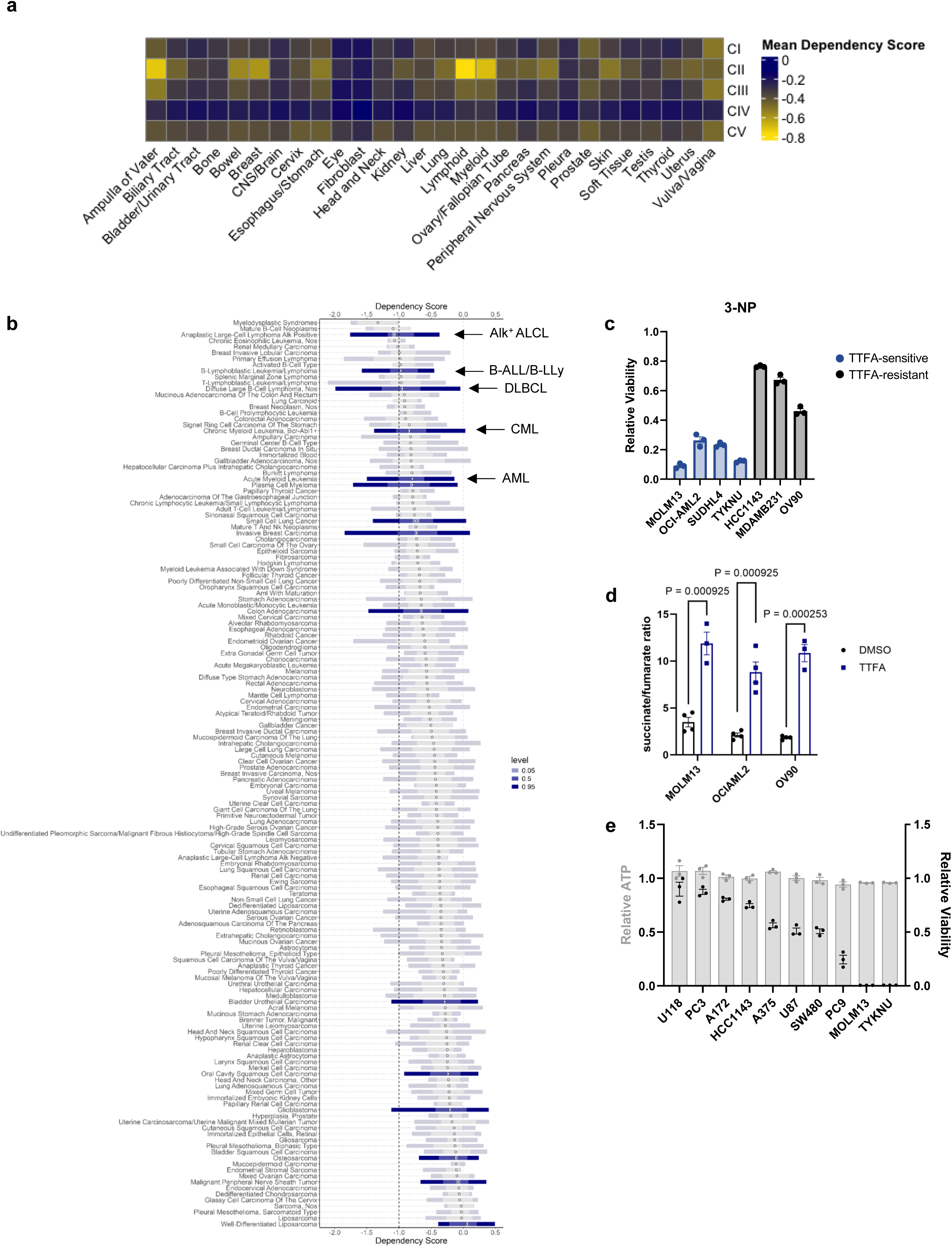
a) Heatmap of Dependency scores for each ETC complex across all cancer lineages represented in the DepMap database. Dependency scores are expressed as the mean score for core and accessory ETC genes within a given lineage, with a more negative number indicating a stronger dependency. b) Complex II sub-lineage dependencies in cancer. Horizontal bars represent ranges of dependency scores for cancer sub-lineages, with differential Complex II dependency between sub-lineages indicated by solid blue versus transparent gray bars. Shades of blue or gray within each bar represent the proportion of cells with a given dependency score interval, annotating 5%, 50% and 95% of cells within each sub-lineage (indicated in the legend as ‘level’). Leukemia and lymphoma subtypes with dependency scores that are significantly different from other sub-lineages are shown (Alk^+^ ALCL, Alk^+^ anaplastic large cell lymphoma; B-ALL/B-LLy, B-cell acute lymphoblastic leukemia/B-cell lymphoblastic lymphoma; DLBCL, diffuse large B-cell lymphoma; CML, chronic myeloid leukemia; AML, acute myeloid leukemia). c) Effect of Complex II inhibitor 3NP (2 mM) on viability of select cell lines that are either sensitive or resistant to TTFA. d) The effect of Complex II inhibition on the succinate:fumarate ratio of cell lines sensitive (MOLM-13, OCI-AML2) or resistant (OV-90) to TTFA. e) Differential effect of Complex II inhibition by TTFA on ATP levels (grey bars) assayed after 30 minutes compared to viability (black squares) recorded after 3 days in culture with TTFA.

**Extended Data 3 (Related to Fig. 2).**
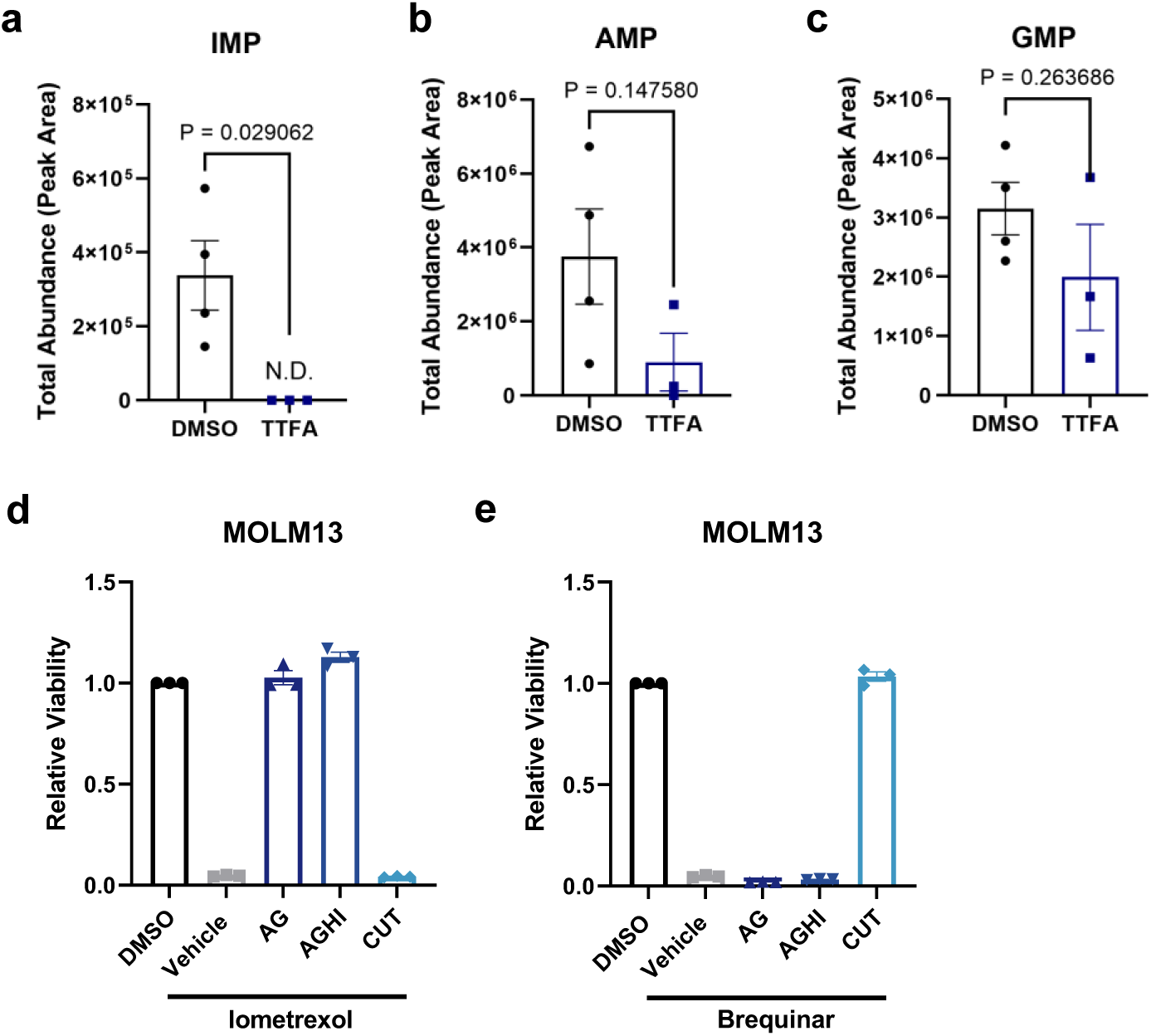
a-c) Effect of Complex II inhibition by TTFA on levels of *de novo* purines (a) IMP, (b) AMP/dGMP, and (c) GMP. AMP/dGMP cannot be resolved based on mass spectrometry separation. d) Effect of exogenous purines (adenine (A), guanine (G), hypoxanthine (H), inosine (I)) or pyrimidines (cytidine (C), uridine (U), thymidine (T)) on viability in MOLM-13 cells treated with the targeted purine synthesis (GART) inhibitor, lometrexol (LTX). Each nucleotide species was added back at a concentration of 30 μM. e) Effect of exogenous purines or pyrimidines on viability in MOLM-13 cells treated with dihydroorotate dehydrogenase (DHODH) inhibitor, brequinar (2 μM). f-h) Data were analyzed using one-way ANOVA with a Tukey post-hoc test.

**Extended Data 4 (Related to Fig. 3).**
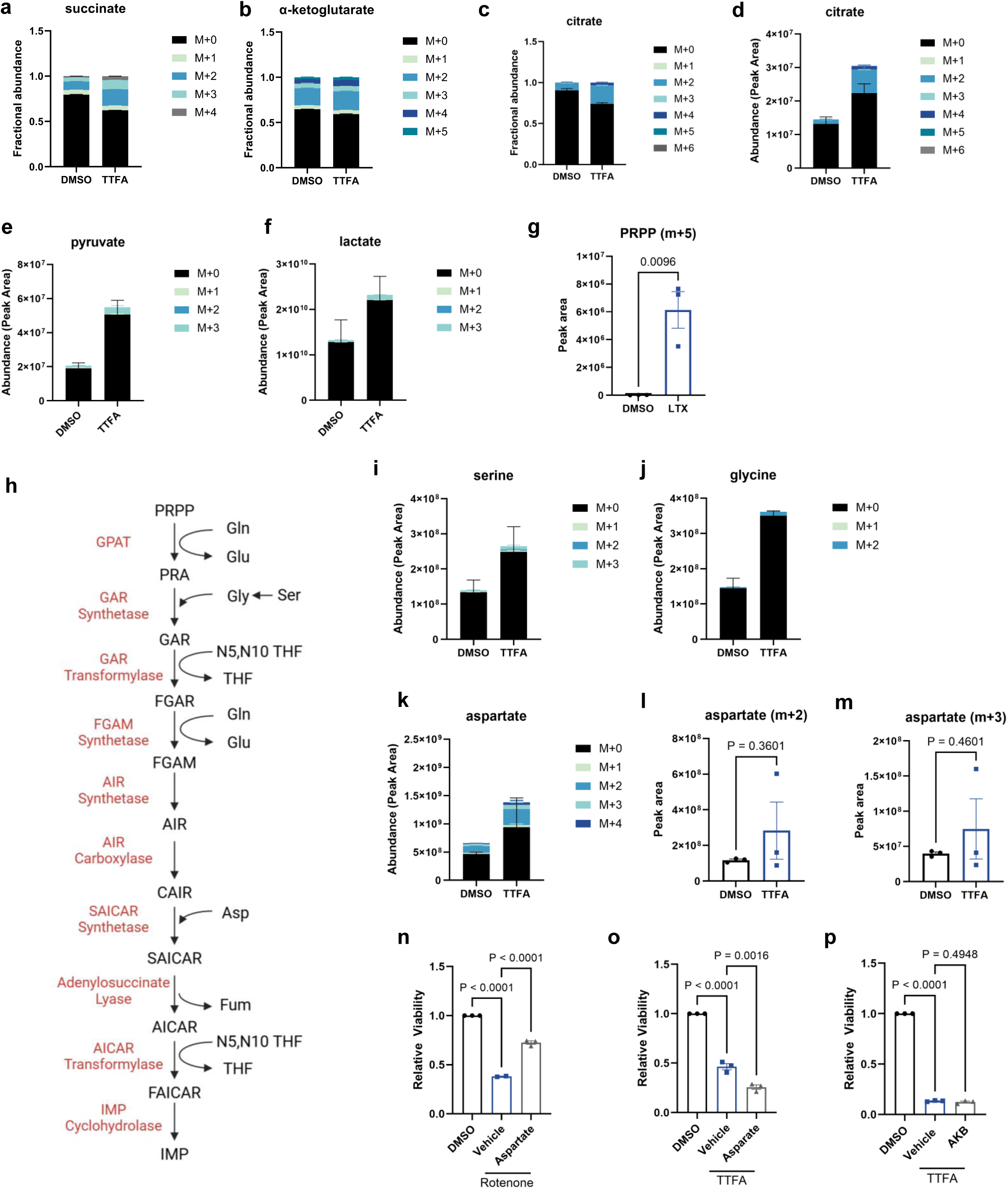
a-c) Effect of Complex II inhibition on the fractional composition of (a) succinate, (b) α-ketoglutarate, and (c) citrate from ^13^C_6_-glucose tracing studies. d) Isotope peak areas for citrate with and without Complex II inhibition. e-f) Effect of Complex II inhibition on ^13^C_6_-glucose-derived isotope content of glycolytic metabolites (e) pyruvate and (f) lactate. g) Quantification of ^13^C_6_-glucose derived m+5 PRPP after culture in the presence or absence of lometrexol (LTX). h) Overview of the *de novo* purine biosynthesis pathway, including contributions from serine (Ser), glycine (Gly), ethylene tetrahydrofolate (N5, N10 THF), glutamine (Gln), and aspartate (Asp). Glutamate (Glu), tetrahydrofolate (THF) and fumarate (Fum) are reaction products. Metabolite intermediates are in black, enzymes in red. i-j) Effect of Complex II inhibition on the ^13^C_6_-glucose-derived isotope content of (i) serine and (j) glycine. k-m) Effect of Complex II inhibition on ^13^C_6_-glucose derived aspartate isotopes, including (k) total peak area for aspartate, (l) peak area for m+2 aspartate and (m) m+3 aspartate, which are derived from oxidative and reductive pyruvate metabolism, respectively. n) Effect of aspartate add-back (20 mM) on the viability of MOLM-13 cells when cultured in control (DMSO group) conditions or with the Complex I inhibitor, rotenone (Vehicle and Aspartate groups). o-p) Effect of either (o) aspartate or (p) alpha-ketobutyrate (AKB, 1 mM) add-back on the viability of MOLM-13 cells cultured in DMSO or the Complex II inhibitor, TTFA. Data shown are mean ± s.e.m. of 3 replicate experiments. Between group metabolite comparison were made using unpaired t-tests and viability assessments made with one-way ANOVA and Tukey’s post-hoc HSD.

**Extended Data 5 (Related to Fig. 3).**
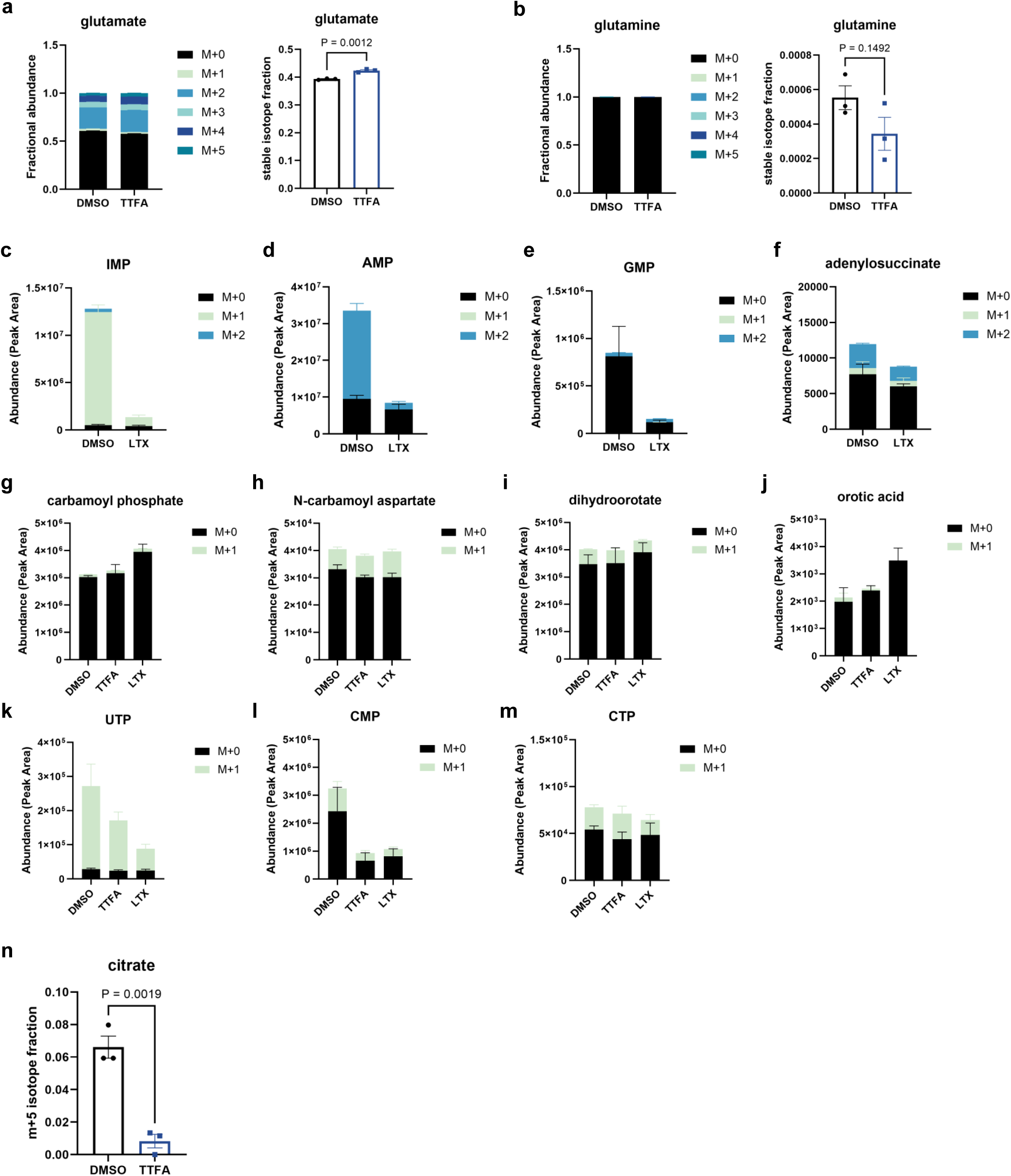
a) Effect of Complex II inhibition on ^13^C_6_-glucose derived isotopes in glutamate. Data at left show the fractional abundance of all glutamate isotopes, data at right show the total isotope fraction before and after Complex II inhibition. b) Effect of Complex II inhibition on ^13^C_6_-glucose-derived glutamine isotopes. Data presented as in panel (a). c-f) Effect of lometrexol (LTX) on the incorporation of ^15^N_1_-glutamine into *de novo* purines (d) IMP, (e) AMP, (f) GMP and (g) adenylosuccinate. g-m) Effect of either Complex II (TTFA) or LTX on ^15^N_1_-glutamine derived nitrogen into pyrimidines. Abbreviations: UTP, uridine triphosphate; CMP, cytidine monophosphate; CTP, cytidine triphosphate. n) Effect of Complex II inhibition on the fractional content of m+5 citrate derived from ^13^C_5_-glutamine. All data are shown as mean ± s.e.m. of 3 independent experiments. Between group comparisons were performed for individual isotopes using an unpaired t-test.

**Extended Data 6 (Related to Fig. 4).**
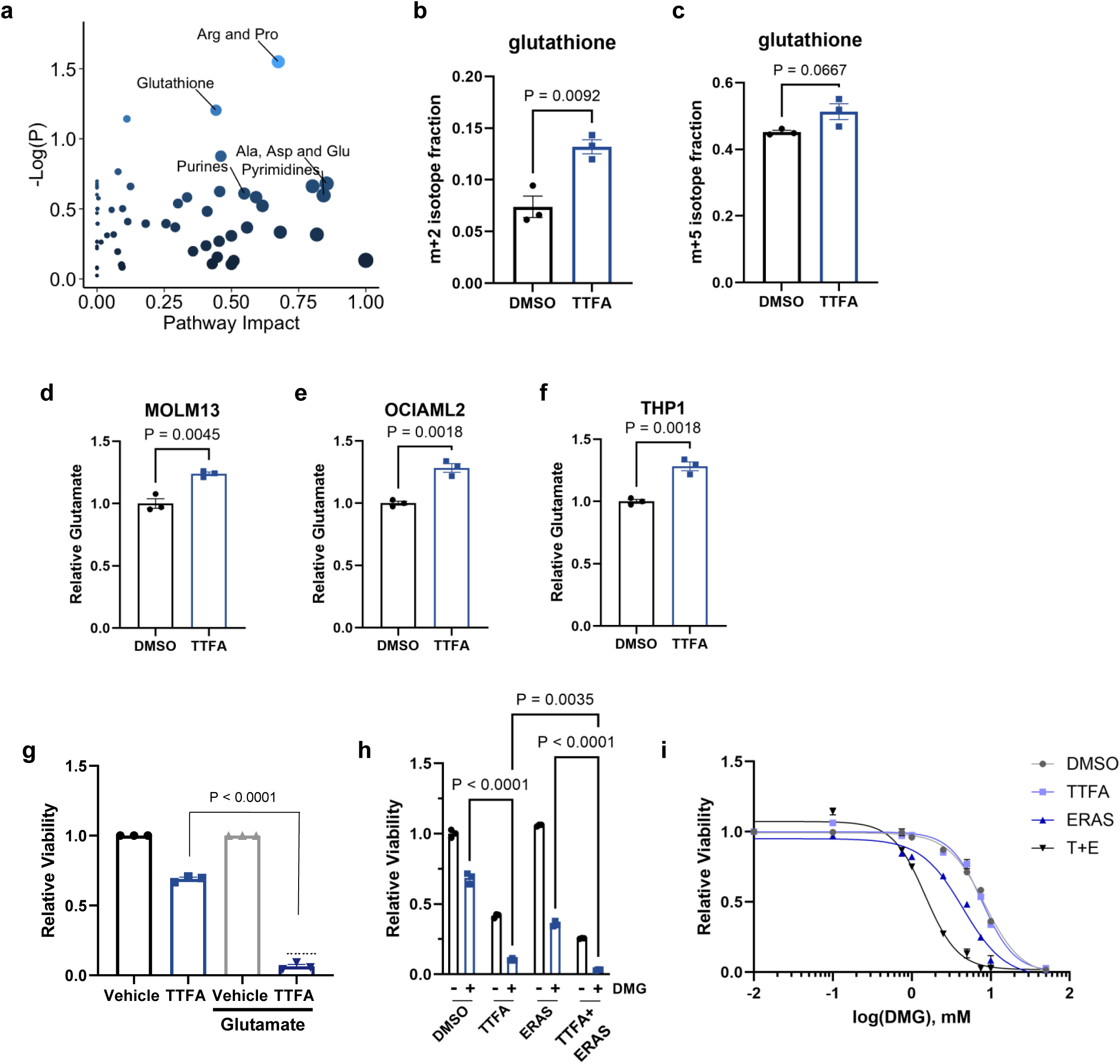
a) Pathway Impact plot for steady state metabolites in MOLM-13 cells treated with DMSO or TTFA for 18hr. Pathway Impact data and p-values were calculated using Metaboanalyst software (see Extended Data Supplemental Table 2). b-c) Effect of Complex II inhibition on fractional composition of glutathione (GSH) from either (b) ^13^C_6_-glucose (m+2 GSH, from either glutamate or glycine) or (c) ^13^C_5_-glutamine (m+5 GSH, from glutamate). d-f) Relative glutamate levels after 24hr TTFA treatment in (d) MOLM-13, (e) OCI-AML2 or (f) THP-1 cells. g) Effect of 5 mM glutamate in addition to 75 μM TTFA on viability after 72hr of culture in OCI-AML2 cells. h) Effect of TTFA (200 μM) and/or erastin (2.5 μM) on the viability of MOLM-13 cells treated in the presence of absence of DMG for 48hr. Concentrations of TTFA and erastin are the same as in Fig. 4b. i) Effect of TTFA (T, 100 mM) and/or erastin (E, 2.5 mM) on the sensitivity of MOLM-13 cells to increasing concentrations of TTFA. GI50 values were calculated from these replicates and presented in Fig 4c. All data are shown as mean ± s.e.m. of 3 independent experiments. For panels b-f an unpaired, two-tailed Student’s t-test was used to determine between group differences. In panel g, an ANOVA was used with Tukey post-hoc, and for panel h, a two-way ANOVA was used (drug treatment x DMG) followed by testing multiple comparisons.

**Extended Data 7 (Related to Fig. 5).**
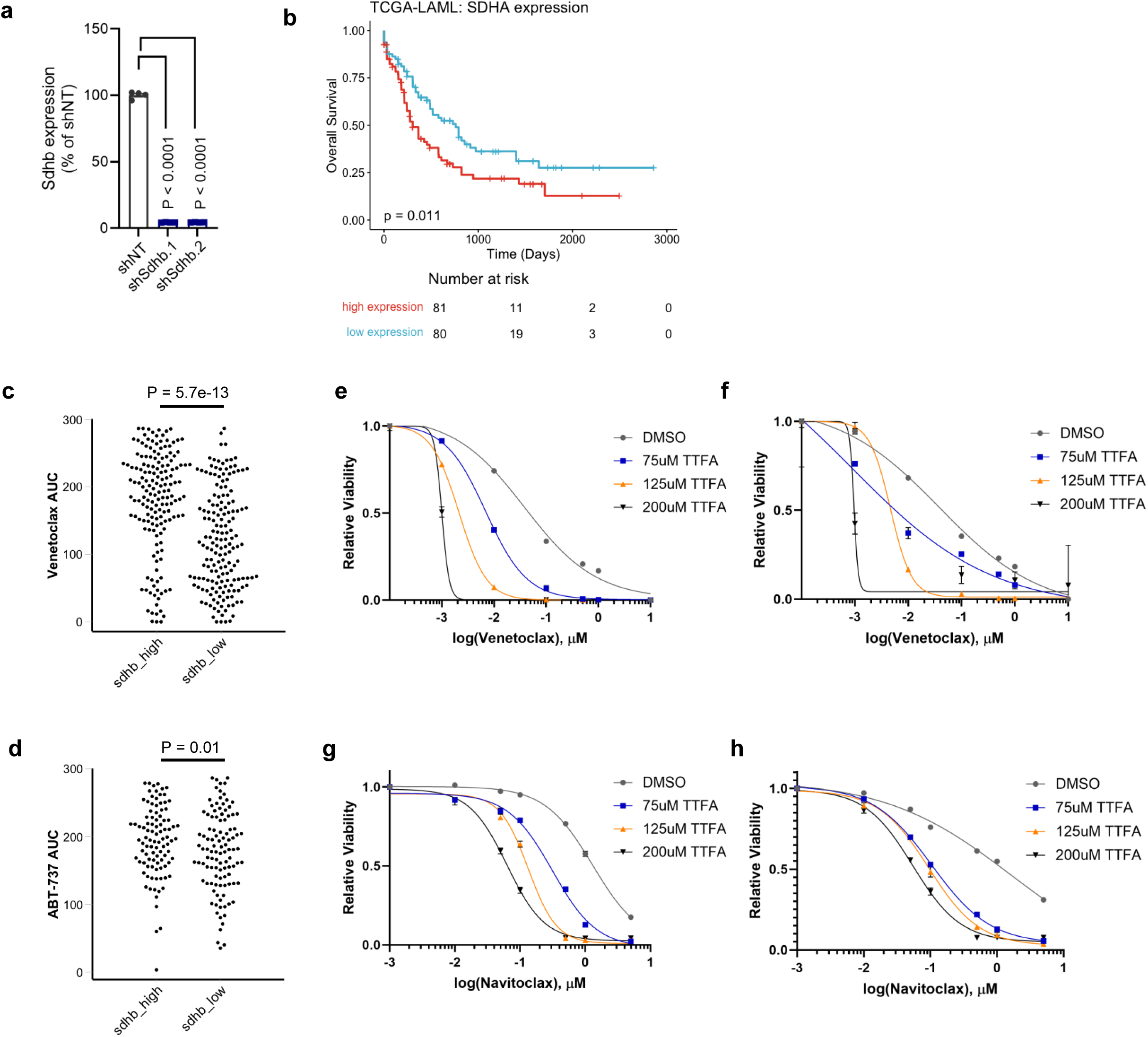
a) *Sdhb* gene expression by qPCR in MLL-AF9 cells 5 after doxycycline induction of non-targeting (shNT) or *Sdhb*-targeting (shSdhb.1 or shSdhb.2) small hairpin RNA. b) Overall survival in the TCGA-LAML cohort based on *SDHA* expression above (high expression, red) or below (low expression, blue) the mean of the entire cohort. c) Beat AML area under the curve (AUC) data for venetoclax treatment, stratified by *SDHB* levels above (sdhb_high) or below (sdhb_low) the median. d) Beat AML AUC data for BCL-2 inhibitor ABT-737, stratified by *SDHB* levels above (sdhb_high) or below (sdhb_low) the median. e-f) Effect of Complex II inhibition with increasing doses of TTFA on AML sensitivity to venetoclax in (e) MOLM-13 or (f) OCI-AML2 cells. g-h) Effect of Complex II inhibition with increasing doses of TTFA on AML sensitivity to navitoclax in (g) MOLM-13 and (h) OCI-AML2 cells.

## METHODS

### Data Download

Gene effect data from Project Achilles were downloaded from the DepMap portal at: depmap.org.

### Data Availability

All data associated with this study are available in the main article or extended data. The Broad Institute has information on their website and the website dedicated to the Dependency Map project about how the raw data were generated and provides a list of references. Results from the pathway-pathway coessentiality are provided as a web application at www.datadrivenhypothesis.org, which was developed by the Hirschey Lab. Code is available on GitHub (https://github.com/hirscheylab/ddh) for a package we generated that contains functions to create plots and tables, and https://github.com/hirscheylab/ddh_app for the web app).

### Code Availability

All code is available on GitHub (https://github.com/hirscheylab/ddh).

### Gene Ontology Embedding and Canonical Correlation Analysis

To develop a pathway-pathway coessentiality map, we started with essentiality data from the Cancer Dependency Map. Gene sets representing pathways were identified using the MSigDB resource. Next, Principal Components Analysis (PCA) was performed independently for each pathway. The first four principal components for each pathway were selected, reducing all pathways from their original respective sizes to a common size of four components. This dimension reduction was performed to make the pathways comparable and reduce overfitting of the Canonical Correlation Analysis (CCA) model, which can occur when the number of variables is high and large differences exist between the number of variables in compared gene sets. Finally, pathway-level co-dependency data were extracted by computing CCA on the first four Principal Components (PCs), the results of which were then used to select the first canonical correlation (CC1). CC1 values were used to represent pathway coessentiality with a range of 0-1.

For ETC embeddings, custom pathways were created using core and accessory ETC gene subunits. Genes for each complex were selected as annotated in Mitocarta 3.0. The number of genes used for each ETC are as follows: Complex I, 37 genes; Complex II, 4 genes; Complex III, 10 genes; Complex IV, 25 genes; Complex V, 18 genes. PCA analysis, extraction of top PCs, and CC1 calculation were performed identically to the pathway analysis described above.

To determine how CCA changed correlation scores relative to simple coessentiality correlations, we extracted dependency scores for ETC genes from DepMap used them to calculate coessentiality between each ETC gene and each GO pathway. These single gene-pathway correlations were combined to represent ETC complex-pathway correlations and displayed as ‘Gene-Pathway’ values. These were compared to canonical correlation scores that were calculated using CCA, which involved first extracting the principal components for ETC complexes, as described above. These values are represented as ‘Complex-Pathway’.

To compare Complex II dependency scores between cancer sub-lineages,, genes were grouped by ETC complex. The mean dependency score was calculated within a lineage or sub-lineage, as well as the 5%, 50% and 95% intervals. Differences in ETC complex dependency between sub-lineages was calculated using a two-tailed t-test between each lineage and the aggregate of cells not in the lineage.

### Usage

https://www.datadrivenhypothesis.org

### Cell lines and reagents

All cell lines were purchased from the American Type Culture Collection (ATCC) or Duke University Cell Culture Facility (CCF) and maintained at 37°C with 5% CO_2_. All cell lines were maintained in RPMI 1640 medium with 10% fetal bovine serum (FBS) and 1% penicillin-streptomycin (P/S). For experiments with addition of exogenous metabolites and glutamate modulation experiments, RPMI 1640 medium with 10% dialyzed FBS and 1% P/S was used. TTFA and AA5 were purchased from Cayman Chemical. Lometrexol, 3-NP, and all exogenous metabolites were purchased from Sigma Aldrich, except for cytosine and thymine, which were purchased from TargetMol. Brequinar was purchased from MedChem Express. Erastin was purchased from Apexbio. Dimethyl-glutamate was purchased from Sigma. Venetoclax and navitoclax were purchased from Selleck chem.

### Cellular viability assays

Cells were seeded at an equal density ranging from 2-10e^3^ cells per cm^2^ in tissue culture plates and allowed to equilibrate. After 24 hours, treatment drugs or metabolites were added to cells in triplicate. Cell viability was measured 48-96 hours later using Cell-Titer Glo (Promega) 1:1 v/v on a Tecan plate reader (Infinite M1000 PRO). The drug and metabolite concentrations, as well as the specific timing of each experiment, are noted in figure legends. Relative cell viability was calculated by normalizing treatment luminescence values to vehicle-treated wells. For experiments with drug or with metabolite combinations, the second drug or metabolite was kept at a constant concentration across all wells and the first drug was applied on top of the background drug or metabolite(s). Relative cell viability was then normalized to the luminescence of cells in the presence of the background drug or metabolite alone.

### Measurement of ATP levels

Cells were seeded at a density of 10e^3^ cells per well in a 96-well plate. After 24 hours, cells were treated with DMSO or TTFA (200μM) for 30 minutes in triplicate, before viability was impacted, and assayed using Cell-Titer Glo (Promega) on a Tecan plate reader (Infinite M1000 PRO). Luminescence values were normalized to DMSO-only conditions to obtain relative ATP levels.

### Metabolite profiling and stable isotope tracing

For steady-state metabolite profiling, 1e^6^ cells were seeded in 10 cm plates. After 24 hours, DMSO or TTFA (200 μM) was added to cells in triplicate. After 18 hours of treatment, cells were harvested by centrifugation, washed with ice-cold 0.9% saline twice, and 80% (vol/vol) methanol pre-chilled to -80°C was added to cells on dry ice to quench and extract metabolites. Lysates were incubated at -80°C for 5 minutes, brought to room temperature for 15 minutes, and then vortexed for 1 minute. The freeze-thaw steps were repeated three times, after which samples were incubated overnight at -20°C to precipitate proteins. Samples were vortexed for 30 seconds and centrifuged for 15 minutes at 20,000x*g* at 4°C. Supernatants were stored at -80°C until analysis by LC-MS. Samples were normalized to protein concentration contained in the cell pellet. Briefly, cell pellets were solubilized in 4M urea and protein concentration was measured using the Bradford assay.

For stable-isotope tracing experiments, 1e^6^ cells were seeded in 10 cm plates. After 24 hours, cell culture media was changed to glucose-free of glutamine-free media containing 10% dialyzed FBS and either 11.1 mM ^13^C_6_-glucose, 2 mM ^15^N_1_-glutamine-amide or ^13^C_5_-glutamine (Cambridge Isotope Laboratories), respectively. Simultaneously, vehicle, TTFA (200 μM), or LTX (150 nM) was added for 24 hours. Metabolite extraction was performed as above. Samples were normalized to cell count.

For quantitative nucleotide metabolomics experiments, 30e^6^ cells were seeded in 15 cm plates with 10% dialyzed FBS as above. After 24 hours, cells were treated with 5 mM DMG for 18 hours. Cells were harvested into pre-chilled 100% MeOH and flash frozen in liquid nitrogen for subsequent metabolite extraction.

For all metabolomics experiments, samples were reconstituted after extraction by first drying the extraction solution using a SpeedVac and 60% acetonitrile was added to the tube, followed by vortexing for 30 seconds. Reconstituted samples were then centrifuged for 30 minutes at 20,000*g* at 4 °C. Supernatant was collected for LC-MS analysis.

Samples were analyzed by High-Performance Liquid Chromatography and High-Resolution Mass Spectrometry and Tandem Mass Spectrometry (HPLC-MS/MS). Specifically, the system consisted of a Thermo Q-Exactive in line with an electrospray source and an Ultimate3000 (Thermo) series HPLC consisting of a binary pump, degasser, and auto-sampler outfitted with a Xbridge Amide column (Waters; dimensions of 3.0 mm × 100 mm and a 3.5 µm particle size). The mobile phase A contained 95% (vol/vol) water, 5% (vol/vol) acetonitrile, 10 mM ammonium hydroxide, 10 mM ammonium acetate, pH = 9.0; B was 100% Acetonitrile. The gradient was as follows: 0 min, 15% A; 2.5 min, 30% A; 7 min, 43% A; 16 min, 62% A; 16.1-18 min, 75% A; 18-25 min, 15% A with a flow rate of 150 μL/min. The capillary of the ESI source was set to 275 °C, with sheath gas at 35 arbitrary units, auxiliary gas at 5 arbitrary units and the spray voltage at 4.0 kV. In positive/negative polarity switching mode, an *m*/*z* scan range from 60 to 900 was chosen and MS1 data was collected at a resolution of 70,000. The automatic gain control (AGC) target was set at 1 × 10^6^ and the maximum injection time was 200 ms. The top 5 precursor ions were subsequently fragmented, in a data-dependent manner, using the higher energy collisional dissociation (HCD) cell set to 30% normalized collision energy in MS2 at a resolution power of 17,500. Besides matching m/z, metabolites are identified by matching either retention time with analytical standards and/or MS2 fragmentation pattern. Data acquisition and analysis were carried out by Xcalibur 4.1 software and Tracefinder 4.1 software, respectively (both from Thermo Fisher Scientific). For isotope tracing experiments, natural isotope abundance was corrected for using AccuCor.

### Glutamate assay

Cells were seeded at 0.5-1e^5^ per cm^2^ in culture plates. After 24 hours, DMSO or given drug or metabolite was added to cells in triplicate for an additional 24-48 hours. At the end of the culture period, an aliquot of cells from the cultured plates was used to calculate viability with the Cell Titer Glo assay. Remaining cells were harvested by centrifugation and washed twice with ice-cold PBS. 0.6N HCl was added for 5 minutes to lyse cells and inactivate metabolism. An equivalent volume of 1mM Tris Base was then added to neutralize lysates. The Glutamate-Glo assay (Promega) was performed according to the manufacturer’s instructions to determine glutamate levels. Glutamate levels were normalized to cell viability for each sample and then normalized to vehicle-only samples to obtain relative glutamate levels.

### MLL-AF9 mouse model

Experiments involving mice were conducted in accordance with approved animal use and care protocols. Mice were maintained on a 12:12hr light:dark cycle with *ad libitum* access to food and water. For experiments examining the effect of Complex II in AML *in vivo*, Granulocyte-Macrophage Progenitor (GMP) cells expressing dsRed in tandem with the MLL-AF9 leukemia translocation were isolated and transfected with lentivirus constitutively expressing Venus YFP and a doxycycline-inducible Crimson far-red RFP in tandem with either non-targeting shRNA (shNT) or one of two independent shRNA hairpins targeting Complex II subunit *Sdhb* (shSdhb.1 and shSdhb.2, respectively). The effect of doxycycline on *Sdhb* expression in shNT, shSdhb.1, or shSdhb.2 cells was determined in transfected MLL-AF9 cells after 5 days.

Transfected MLL-AF9 cells were transplanted via tail vein injection into syngeneic recipients. Leukemia burden was quantified as the proportion of AML cells expressing the inducible transcript (Crimson^+^Venus^+^) relative to total AML cells (Venus^+^) in bone marrow or spleen. For survival studies, MLL-AF9 transplanted mice were followed after engraftment until they displayed signs of overt leukemia or if moribund.

### TCGA cancer survival data analysis

TCGA survival data were analyzed using the TCGA survival tool (www.tcga-survival.com). For survival analysis by gene expression status, groups were divided into ‘gene high’ and ‘gene low’ groups based on the mean value for the variable being analyzed. Kaplan Meier curves were generated using R software and between group statistics performed with a log-rank test. Overall survival hazard ratios were calculated for each cancer subset using Cox proportional hazard regression analysis.

### Statistical analysis

All data are represented by mean ± standard error of the mean (s.e.m.). P values were calculated using a combination of methods including two-tailed t-tests, analysis of variance, and post-hoc pairwise comparisons as indicated in figure legends. P values in the figures are presented for individual isotope content or for viability assays.

## Acknowledgments

We want to thank: Anne Le, Shenghao Guo, and Pratik Khare at John’s Hopkins University for assistance with pilot isotope tracing experiments and the Metabolomics Core Facility at Robert H. Lurie Comprehensive Cancer Center of Northwestern University for providing metabolomics services; Dan Leer and Hilmar Lapp at Duke University for computational and technical support; Cédric Scherer for data visualization assistance. We would like to acknowledge funding support from the Glenn Foundation (MDH), the National Institutes of Health and the NIA grant R01AG045351 (MDH), NCI grant R01CA266389 (KCW), and the Duke Cancer Institute P30 Cancer Center Support Grant P30CA014236 (KCW and MDH). DKZ was supported through Duke’s Department of Pediatrics and funding sources R38AI140297 (NIAID), T32HD094671 (NICHD) and 1K38CA282963 (NCI). AES was supported by F31CA254127 (NCI). The content is solely the responsibility of the authors and does not necessarily represent the official views of the National Institutes of Health or other funding sources.

## Additional Information

Correspondence and requests for materials should be addressed to KCW or MDH.

